# No Brain Signature of Prediction Error to Unattended Visual Stimuli Despite Early Brain Encoding of the Stimuli

**DOI:** 10.1101/2022.10.04.510779

**Authors:** Alie G. Male, Robert P. O’Shea

## Abstract

Prediction error is a basic component of predictive-coding theory of brain processing. According to the theory, each stage of brain processing of sensory information generates a model of the current sensory input; subsequent input is compared against the model and only if there is a mismatch, a prediction error, is further processing performed. Recently, Smout et al. [1] found that a signature of prediction error, the visual (v) mismatch negativity (MMN), for a fundamental property of visual input—its orientation—was absent without attention on the stimuli. This is remarkable because the weight of evidence for MMNs from audition and vision is that they occur without attention. To resolve this discrepancy, we conducted an experiment addressing two alternative explanations for Smout et al.’s finding: that it was from a lack of reproducibility or that participants’ visual systems did not encode the stimuli when attention was on something else.

We conducted a similar experiment to Smout et al.’s. We showed 21 participants sequences of identically oriented Gabor patches, standards, and, unpredictably, otherwise identical, Gabor patches differing in orientation by ±15°, ±30°, and ±60°, deviants. To test whether participants encoded the orientation of the standards, we varied the numbers of standards preceding a deviant, allowing us to search for a decrease in activity with the number of repetitions of standards—repetition suppression. We diverted participants’ attention from the oriented stimuli with a central, letter-detection task.

We reproduced Smout et al.’s finding of no vMMN without attention, strengthening their finding. We also found that our participants showed repetition suppression: they did encode the stimuli pre-attentively. We also found early processing of deviants. We discuss whether this earlier processing of deviants may be why no further processing, in the vMMN time window, occurs.

---

Our senses are flooded by information. For example, the four million cones of the human retina encode a staggering 101,200,000 bits of information at any instant if their responses to light were binary (which they are not) [2]. Still, we do not experience a flood—only a stream—of information to which we pay attention or of which we become conscious [3]. What we see does not require such an impossible burden of encoding because it is highly redundant, allowing prediction of the state of one photoreceptor at any instant from its state the instant before and from the state of its neighbors. This seminal idea developed into predictive-coding theory, first of the retina [4] and then of the brain’s hierarchical sensory systems [5, 6].

Predictive coding theory is a leading theory of how the brain deals with sensory input [7]. It is that the brain uses recent, past, bottom-up, sensory information, along with top-down information (such as prior probabilities, expectations, and attention) to generate predictive models of sensory input at various levels of the sensory pathway. If future inputs match the prediction, no further processing is required. If they do not match—a so-called prediction error—further processing occurs to update the model.

We studied the visual mismatch negativity (vMMN) as our signature of prediction error [8]. The vMMN is an event-related potential (ERP) from electroencephalography. Stefanics et al. [8] have reviewed its hundreds of studies: It is evoked by an unpredicted change in what we see. It occurs when a rare, unpredicted, deviant, visual stimulus occurs after a sequence of identical, standard, visual stimuli in a so-called oddball sequence. It is greater ERP negativity from parieto-occipital (PO) recording sites for deviants than for standards, occurring between 150 ms and 300 ms after the onset of the deviant. It occurs without adaptation, showing the genuine vMMN [9]. It is supposed to occur without attention on the deviant property of the stimuli.

Recently however, Smout et al. [1] found no ERP evidence of the genuine vMMN without attention to unexpected changes in orientation. In their attended condition, they showed each participant displays consisting of an annular Gabor patch with a central black spot. They asked the participant to look only at the spot but to pay attention to the bars of the Gabor and to press a button on rare occasions when the bars’ spatial frequency increased. In their ignored condition, the same participant viewed the same displays but paid attention to the spot and pressed a button on rare occasions when its contrast decreased. The difficulty of the two tasks was equated for each participant in prior psychophysical testing.

Smout et al.’s stimuli of interest were the Gabor annuli. Participants viewed different-length sequences of displays of identical annuli—standards— followed by an otherwise identical annulus but with bars differing in orientation by ±20°, ±40°, ±60°, or ±80°—a deviant. On the next display, the deviant annulus was repeated, commencing a new sequence for which it now served as the standard—the so-called roving-standard paradigm [10]. To measure the genuine vMMN, Smout et al. also showed control sequences in which all possible orientations occurred in random order, from which no expectation could be established, yet which had the same overall frequency of stimuli identical to deviants, equating adaptation—the equiprobable control [11].

Smout et al. found a genuine vMMN from PO electrodes to unexpected changes in the orientation of the bars in the attended condition (between about 170 ms to about 300 ms), but not in the ignored condition (their Figure S1B). They did find genuine, deviant-related responses in the ignored condition: a negativity from about 330-430 ms from a midline central electrode (Cz) and from about 300-470 ms from a midline frontal electrode (Fz). But these are too late to be considered vMMNs, or any sort of prediction error from such an early brain-processing feature of visual input as orientation [12].

Smout et al. confined their remaining clever analyses to their attended condition to answer questions about the timing and orientation tuning of prediction error. But their failure to find a genuine vMMN without attention was wildly unexpected. Smout et al. did not discuss it except to cite one study for a similar finding in the auditory modality [13]. To resolve this discrepancy, we conducted an experiment addressing two alternative explanations for Smout et al.’s finding: that it was from a lack of reproducibility or that participants’ visual systems did not encode the stimuli when attention was on something else.

We made some changes to improve our study’s ability to detect the genuine vMMN and to enhance its design. Smout et al.’s stimuli were Gabor annuli with an outer diameter of 11° diameter and an inner diameter of 0.83°; their inner grey patch interrupted the central bars over a region of about the size of the human foveola to which about 20% of the processing of the visual system is devoted [14]. To improve our chances of finding a vMMN, our Gabor patches were double the size of Smout et al.’s outer diameter size, and our central bars were continuous except for a very small area occupied by the lines of small task-relevant fixation letters. We also omitted an attended condition, eliminating any possible confound in Smout et al.’s design from participants’ moving their eyes to the bars [15], thus bringing them onto central vision. We also analyzed clusters of electrodes around the single electrodes Smout et al. reported. This was to avoid issues related to variability in placement of single electrodes [16].

To look for evidence that participants encoded the orientation of the Gabor patches, we searched for repetition suppression: attenuated brain activity to repeated stimuli [17]. For Gabor patches similar to those used by Smout et al. and by us, this attenuation appears in various components of ERPs, as late as the P2 [17] and as early as the P1 at occipital-parietal electrodes [18].

We found no genuine vMMN to ignored orientation deviants despite showing that participants encoded the orientated stimuli.

## Results

Participants looked at a central, desynchronized stream of letters and pressed a key whenever an X appeared (Figure 1A–B). Desynchronized from the letter stream, Gabor patches appeared with, in experimental blocks, occasional changes in orientation after varying numbers of repetitions of the same orientation and, in control blocks, randomly selected orientations on each trial (Figure 1C–D).

**Figure 1.**
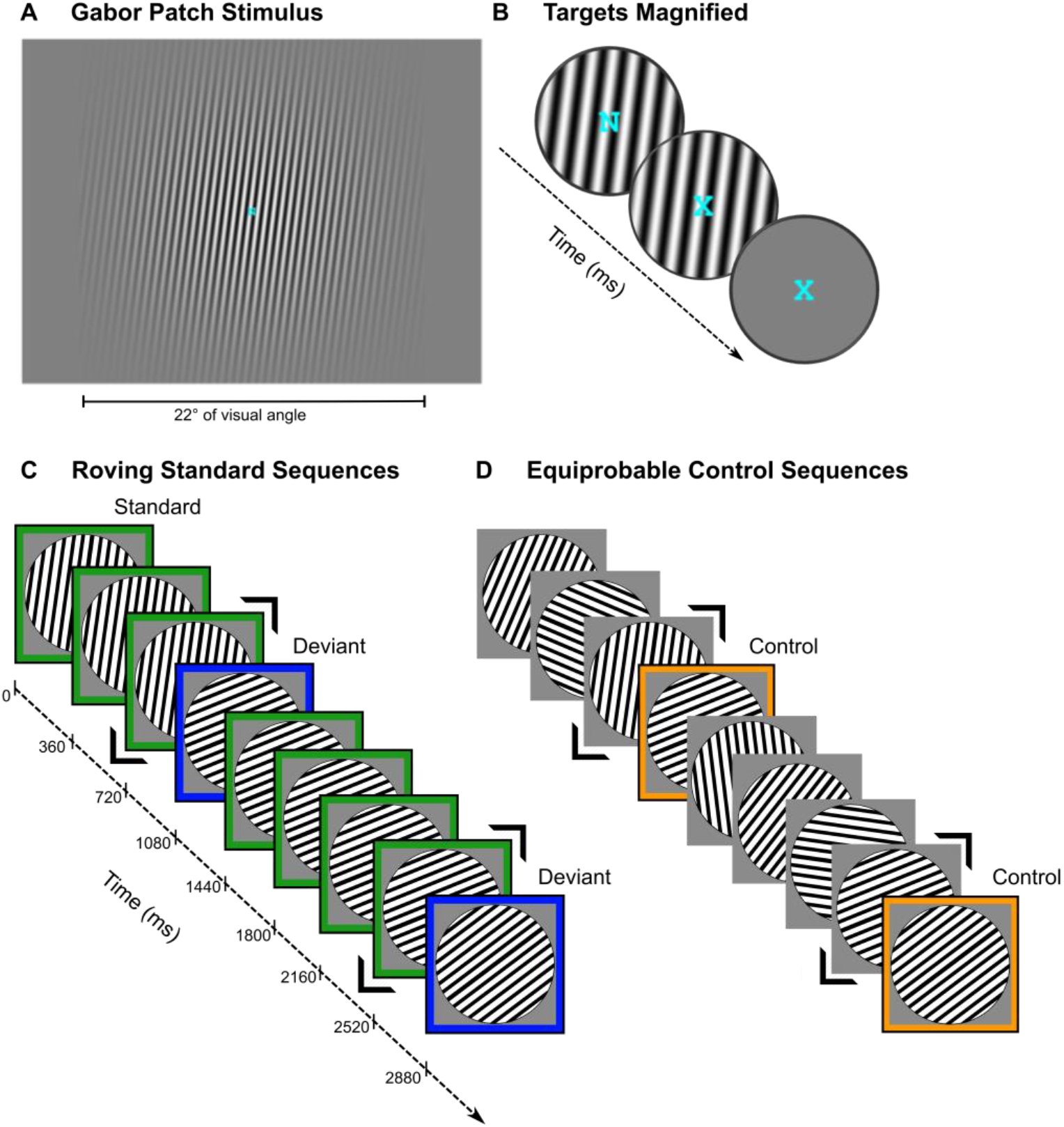
Experimental Paradigm. **A. Screenshots from the experiment of a Gabor patch** tilted 8° clockwise from vertical (0°), with a superimposed fixation letter. **B. Magnified views of a sequence of target letters** superimposed on the Gabor patch (first two images) or on the gray screen during the inter-stimulus interval (ISI). The participants’ task was to press a key whenever an X appeared. Each letter was shown for 600 ms. **C. Illustration of roving standard sequences** in which the first few trials of one sequence is 8°… 8°… 8°… *68*° (i.e., +60° orientation deviant for the first sequence) then …68°… 68°… 68°… 68°… *53°…* (i.e., −15° orientation deviant for the second sequence). Standards are outlined in green; deviants are outlined in blue. The stimulus preceding a deviant is framed with two black chevrons. **D. Illustration of matching equiprobable controls** in which we replaced standard stimuli— except for the ones immediately preceding a deviant from the corresponding roving standard sequence—with Gabor patches whose orientation was randomly chosen among the 12 possible orientations. All orientations appeared equally (8.3%) often. The stimulus in the sequence (also framed with two black chevrons) preceding the one that would have been the deviant in the corresponding roving standard sequence had an identical orientation to that from the corresponding roving standard sequence. The control (outlined in orange) also had the same orientation as that of the deviant from the corresponding roving standard sequence. Gabor patch onset is indicated on the timeline. Stimuli appeared for 80 ms, followed by a grey interstimulus interval (ISI) screen for 280 ms (not shown).

### Did our Participants Attend to the Letter-Detection Task, Thereby Ignoring the Gabor Patches?

We checked whether our participants attended to the central, letter-detection task, thereby ignoring the Gabor patches. The mean hit rate for detecting the target X in the central letter stream was 98.2% and the mean false alarm rate was <0.01%, showing that participants paid attention to the task and did very well on it. There were no significant differences in performance of the task on oddball and control blocks (details of analyses in supplementary materials: S1).

### Did we Find a vMMN to Unpredicted Orientation Changes?

We did not find a genuine vMMN to unpredicted ignored orientation changes. There is no hint of any genuine negativity in the vMMN time range to any unpredicted orientation changes from any electrode cluster (we give details of our analyses, including Bayesian statistical tests, in Table S1). We show the ERPs and the difference waves in Figure 2, formatted similarly to those of Smout et al.’s. For simplicity, we give our results only for the ±60° orientation change, 10° larger than the average orientation change Smout et al. used. We give the ERPs and the difference waves for our other two orientation changes in Figures S1 and S2. Those difference waves also show no evidence of the vMMN. That is, we replicated Smout et al.’s unexpected failure to find a vMMN, strengthening their finding (details in supplementary materials: S2.1). To take another approach to describing our data, we used temporal principal component analysis (PCA; [19]). We found no component that was consistent with a vMMN.

**Figure 2.**
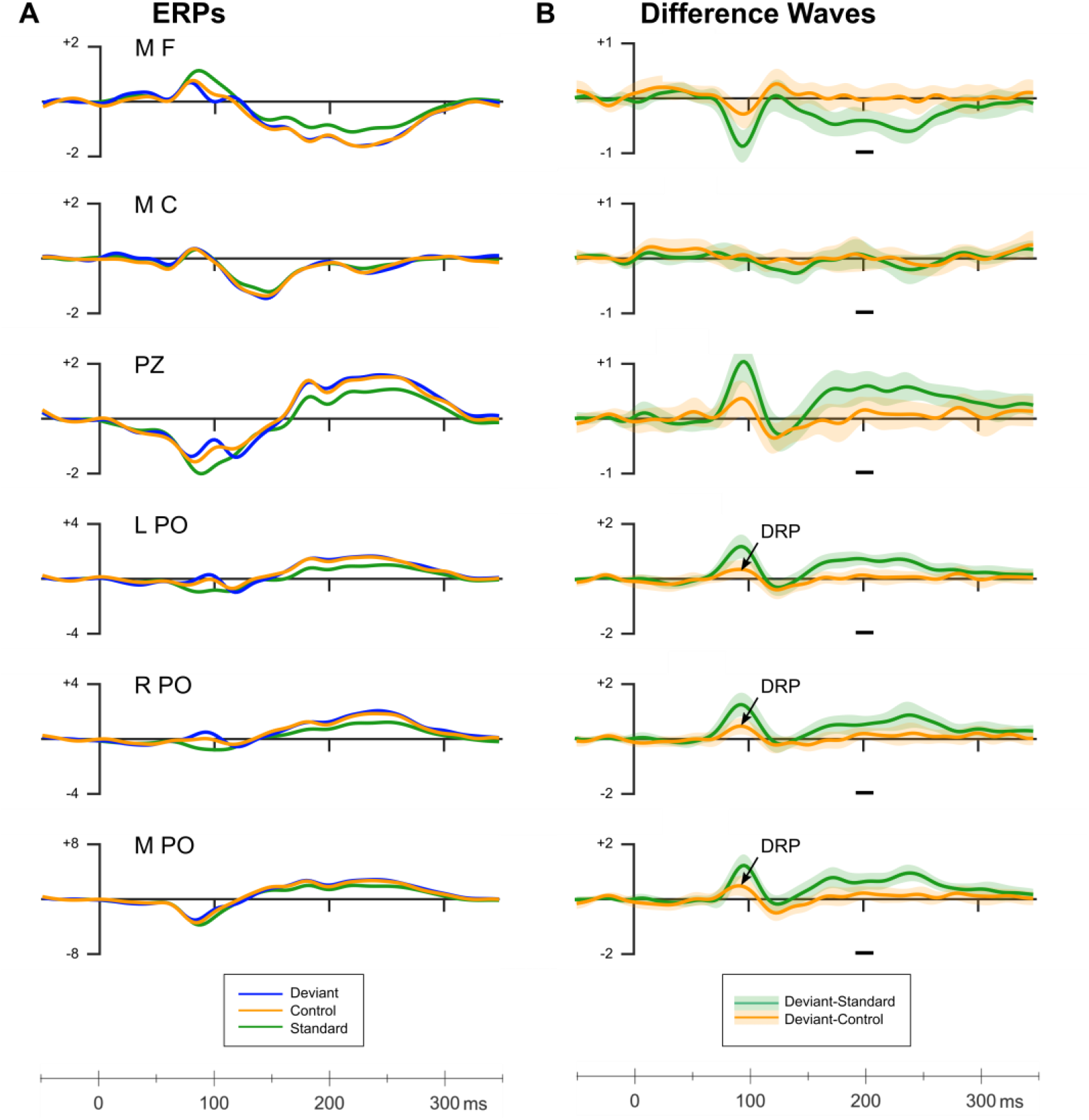
A. **Grand average ERPs from various electrode clusters** around the electrodes Smout et al. reported. B. **Difference waves for classic (60-degree deviant minus standard) and genuine (60-degree deviant minus 60-degree control) deviant-related activity**. The arrowed components show the only genuine deviant-related activity (DRP). Horizontal grey bars illustrate the time window in which Kimura and Takeda (2015) found the largest deviant-minus-control difference (i.e., genuine vMMN) for 32.7° orientation deviants. Amplitudes from this time window were analyzed using Bayesian replication tests (results in Table S1). The lighter colors surrounding the difference waves give ± 1 standard error of the mean.

We also found genuine deviant-related activity outside the vMMN time range: a positivity we call the deviant-related positivity (DRP) from PO electrodes about 90 ms after onset of unpredicted orientation changes. PCA showed that this positivity was from the P1. Figure 3 illustrates these findings (details of analyses in supplementary materials: 2.2).

**Figure 3.**
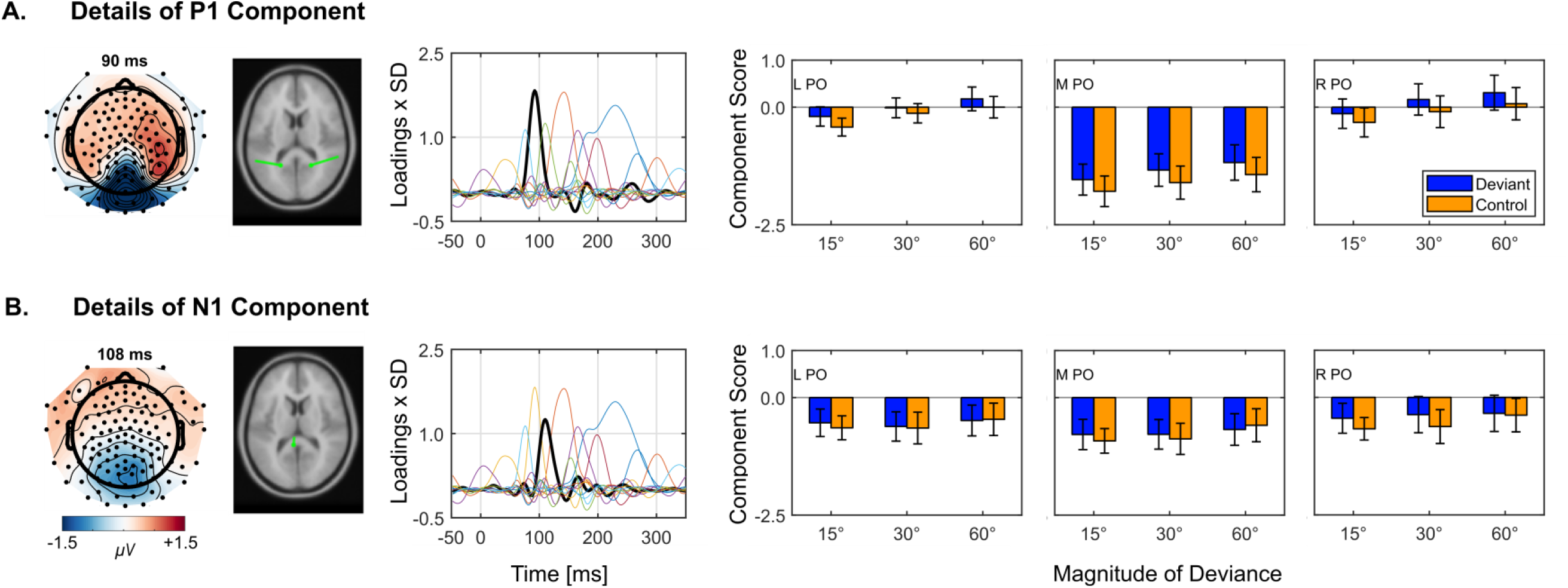
PCA component details for the P1 and N1 components of genuine deviant-related activity. **A. PCA details of the P1. B. PCA details of the N1**. The topographic maps in leftmost column show the combined activity from deviant and control trials at peak latency of the P1 (top) and N1 (bottom). In the second column, we show the results of our dipole analysis of each component. Dipole analysis shows that the pronounced negativity at the M PO reflects the combined negative pole of both P1 components converging within close proximity. In the third column, we show the component loadings (scaled by SD) (thick black line) relative to all other components (thin multi-colored lines). This illustrates the component’s contribution to the overall evoked activity recorded from the scalp. In columns four to six, we show the scores for deviant (orange) and control (purple) trials for each magnitude of deviance (15°, 30°, 60°) at the left (L), midline (M), and right parieto-occipital (PO) region. Error bars depict ±1 standard error. Statistical analyses show that the deviant-related positivity at PO regions in Figure 2 are statistically significant for deviant-minus-control positivity for the P1 only (details in supplementary materials: S2.1).

Smout et al. showed no sign of this early component in their ignored condition. This could be because they obscured the bars of the Gabor patch from central vision by their 0.83°-diameter grey patch, unlike in our study in which the bars crossed central vision.

### Did we Find Repetition Suppression?

We did find repetition suppression (as did Smout et al. in their attended condition). To search for it we used PCA to identify plausible components. We found two components showing repetition suppression—the N1 and P2, most clearly for the P2 (Figure 4). We show the N1 in Figure S3. Figure 4 shows that the PCA P2 scores became significantly less positive with increasing sequential position (i.e., repeated presentations) of standards (green points and regression lines). The P2 scores to random changes (black points and regression lines) became slightly more positive with sequential position, but these are not significant. The results from the standard stimuli show clearly that our participants encoded them, so the absence of a vMMN cannot have been from lack of encoding.

**Figure 4.**
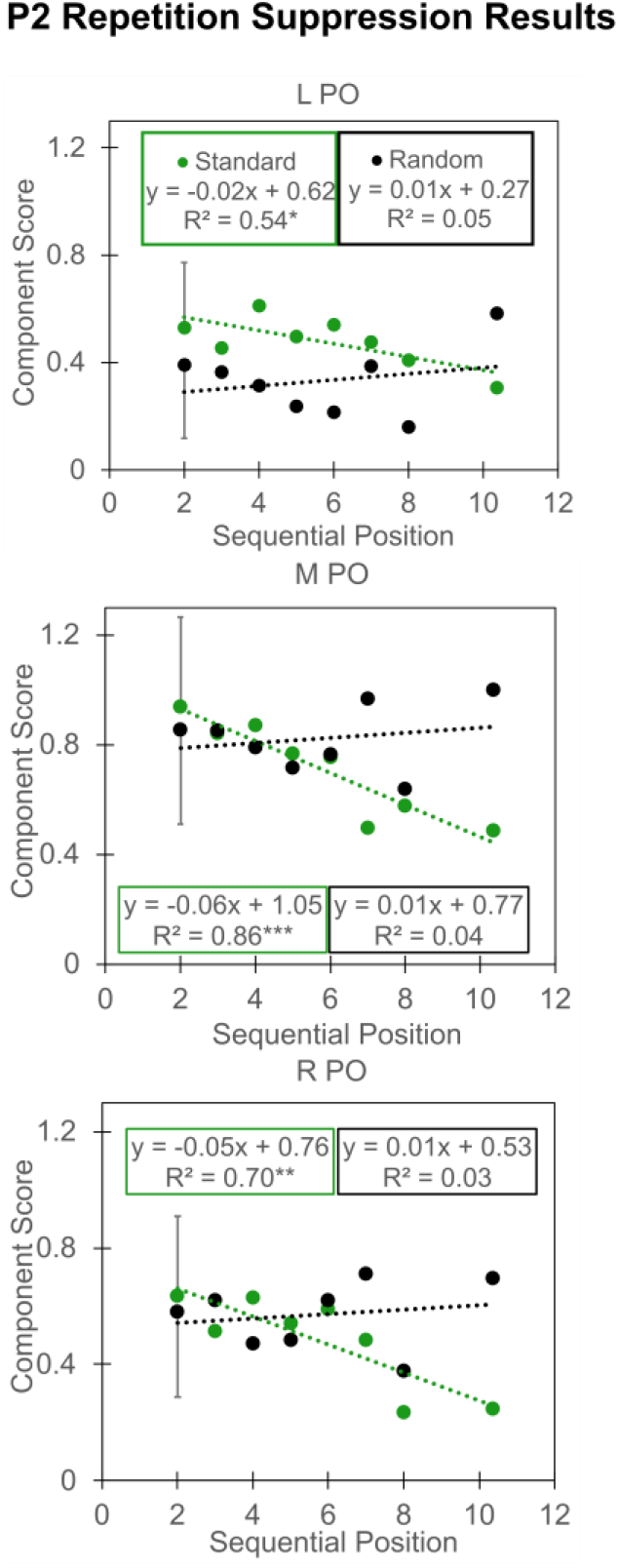
P2 repetition suppression results. P2 PCA scores are shown separately for the left (L), midline (M), and right (R) parietal-occipital (PO) regions of interest (ROI). Regression equations show the dotted lines in the data for standard and random stimuli. Asterisks denote the linear regression significance: \**p*<.05, \*\**p*<.01, *** *p*<.001. Error bars show 1 standard error (SE) calculated as the mean SE for all sequential positions. SE bars appear only on the second sequential position. We show +1 SE for standard and −1 SE for random stimuli.

## Discussion

We have shown that even though participants encoded stimuli, they failed to show a genuine vMMN to unexpected orientation changes when they attended to a central task. Absence of the vMMN is consistent with five experiments including ours [1, 15, 20, 21], but inconsistent with four [11, 22–24]. There are at least three possible explanations for this empirical dilemma:

1. There really is no genuine vMMN to unexpected, unattended orientation changes and the four out of nine studies that reported it are examples of publication bias or methodological errors.
2. There is a genuine vMMN to unexpected, unattended orientation changes and the five studies that failed to find it suffered from some yet-to-be-determined methodological differences or problems.
3. The vMMN is not the appropriate sign of prediction error for isolated and well controlled changes in low-level features of visual input—features that are easily processed in V1.

### 1. No Genuine vMMN to Unexpected Orientation Changes?

Although a 44% hit rate for finding a genuine vMMN is something to take notice of, it is possible there are other negative studies that never made it into print. The vMMN literature does indeed show signs of publication bias. For example, of the 22 orientation vMMN published studies, including those only on the classic vMMN, we are aware of only four reporting no vMMN for a hit rate of 82%. All but one [15] also reported a different condition or experiment in which a vMMN was found, suggesting that vMMN experiments in which all conditions failed to show a vMMN are either not submitted for publication or submitted and rejected. We are aware from a personal communication of at least one such unpublished orientation experiment; there may be more. Such a bias would make it difficult to identify the experimental parameters required to resolve whether the genuine vMMN to orientation changes occurs without attention.

### 2. Methodological Differences or Problems?

We considered whether our experimental parameters were responsible for our results and decided not. Human observers can discriminate orientation differences between gratings of about 0.4° [25], so there is no question that even our smallest orientation difference of 15° is large enough to be perceived. Others have shown genuine vMMNs for other unattended, more complex deviants with presentation times as short as 17 ms [26], much shorter than our 80 ms. And others have shown genuine vMMNs with ISIs as short as 210 ms [27], shorter than our 280 ms. It seems unlikely our choices of orientation change, presentation time, and ISI were responsible for our failure to find a genuine vMMN.

In a review of the literature, we noted that researchers generally failed to isolate individual low-level features, such as orientation, from other, confounding factors including eye movements that are capable of changing simple orientation changes into more complex visual changes [15]. Even Smout et al.’s study suffers from this, with very careful control of fixation in their ignored condition (the participants’ task was to detect contrast changes in a small central spot, requiring them to look at it) but not in their attended condition (the participants’ task was to detect spatial-frequency changes in the bars of the oriented stimuli, 0.42° away from the fixation spot). Male et al. [15] showed evidence that participants’ eyes did stray away from central fixation when the task-relevant information was away from fixation, despite instructions to keep fixation central.

### 3. The vMMN is not the Appropriate Sign of Prediction Error for Low-level Visual Features?

It is possible that unexpected changes in low-level features, such as orientation, are processed earlier than would normally be considered to fall into the time range of the vMMN. We found a genuine, deviant-related positivity at the P1, about 90 ms after onset of the Gabor patch. We have previously reported similar early deviant-related positivity [15] for orientation and for other basic stimulus features. Others have also reported early positivities for deviants, although these do not appear to have been considered as indicators of deviance detection [21, 28–33].

Alternatively, one could argue that we, like Smout et al. and others, did not find the genuine vMMN because prediction error generation in the visual system, unlike the auditory system, requires attention. This would, however, undermine one of the most frequently cited evolutionary purposes of prediction error; to monitor one’s immediate surroundings pre-attentively. Smout et al. carefully navigated this obstacle, saying only that attention enhances the processing of unexpected changes in visual input. We also prefer to leave predictive-coding theory intact and point out that our finding of earlier processing of unexpected orientation changes suggests that no later processing, in the vMMN time range, is necessary.

We hope we have convinced you that the early positivity we found is a useful lead to understanding how the absence of a vMMN without attention could be reconciled with predictive coding theory.

## Methods

We showed 21 [24 in Smout et al.] participants sequences of identically oriented Gabor patches (Figure 1A), standards, and, unpredictably, otherwise identical Gabor patches differing in orientation by ±15°, ±30°, and ±60° [±20°, ±40°, ±60°, or ±80° in Smout et al.], deviants (80 ms duration [100 ms], 280-ms [500 ms] ISI, 634 patches [415] per block; see Figure 1A). Participants looked at a central, desynchronized stream of letters and pressed a key whenever an X appeared (Figure 1B). Standard sequences ranged in length randomly from 3 to more than 11. For an equiprobable control, we presented the same stimuli, but in a random order (see Figure 1D).

### Participants

Twenty-one self-declared healthy adults (8 males, 18 right-handed) with normal or corrected-to-normal vision participated in the experiment (statistical power = .88, based on Kimura and Takeda’s [22] reported effect size for genuine vMMN at PO8: Cohen’s *d* = –0.73). Mean age was 33 years with a range from 18 to 60 years. Most of the participants were undergraduate psychology students at Murdoch University. All participants provided their written informed consent and were free to withdraw from the experiment at any time. Participants received course credit or entry into a draw to win a $50 gift card in return for participation. The Human Research Ethics Committee at Murdoch University approved the experiment (permit 2015 208).

### Apparatus

Participants sat in a light-attenuated chamber viewing a calibrated monitor (17-inch, color, cathode-ray-tube display; Sony Trinitron Multiscan E230) from 57 cm. The monitor showed 1280×1024 pixels (75 Hz refresh rate) and was the only source of light. A chin rest stabilized each participant’s head. Participants gave their responses by pressing a key on a 4-key EGI response box with the index finger of their dominant hand.

A PC running Linux (v4.13.0), GNU Ubuntu (v16.04.4), Octave (v4.0.0) [34], and Psychophysics Toolbox (v3.0.14) [35–37]) delivered the visual stimuli and recorded behavioral responses. An iMac running EGI’s NetStation 5.2 recorded EEG data.

### Stimuli

We used achromatic Gabor patches (mean RGB values of 128 128 128) on an average grey background (RGB values of 128 128 128). A Gabor patch comprises a grating of a particular spatial frequency and orientation whose contrast reduces with distance from the center of the grating according to a Gaussian function [38, 39]. These stimuli are like those used by Smout et al.

Gabor patches had a contrast of .999 [Smout et al. used 1.00; we continue to give their values in brackets after ours], a phase of 0 radians—the black-to-white crossing was in the center of the patch—[not specified, but from their Figure 1, it looks the same], a spatial frequency of 2.4 cycles per degree of visual angle [2.73], and a standard deviation of the Gaussian of 3.84° of visual angle [not specified]. The visible parts of each Gabor patch had a diameter of approximately 22° of visual angle [11°, with a 0.83° central blank area]. There were 12 possible orientations: 8°, 23°, 38°, 53°, 68°, 83°, 98°, 113°, 128°, 143°, 158°, and 173° clockwise from vertical (0°) [nine possible orientations from 0°-160°, in 20° steps].

For the participants’ primary task, capitalized letters were superimposed on the center of the Gabor patch in cyan (RGB values of 0 255 255) in 30-point Courier font [a black fixation spot]. Letters occupied 0.5° (width) × 0.6° (height) with a line width of 0.06° (4 minutes) [visual angle of 0.3° diameter]. We used 25 letters of the English alphabet (all except letter ‘I’). We give an illustration of this in Figure 1.

### Procedure

The participants’ task was to fixate on, and attend to, the continually changing random sequence of letters in the center of the monitor and to press a key with the dominant hand whenever an X appeared. Each letter occurred for 600 ms. None of the letters was immediately repeated. If the participant responded between 0.15 and 1.2 s after target onset, the response was correct. There were 383 letter changes during a block. On average, there were 15 targets (ranging from 7 to 25) per block.

Desynchronized from the letters, we presented our stimuli of interest— concentric Gabor patches—for 80 ms separated by a 280-ms inter-stimulus interval (ISI).

We had two sorts of blocks:

1. Roving standard blocks had 634 Gabor patches of which 114 (18%) were deviants: of different orientation from the immediately preceding Gabor patches. Each deviant became the new standard and repeated until the next deviant. For each participant and block, we organized trials into random-length sequences containing at least three standards (including the deviant of the previous sequence) and no more than 22 standards followed by a deviant. On average, five standards preceded each deviant. The deviant was randomly and equally ±15°, ±30°, or ±60° from the standard. There were 38 trials (6%) for each deviant type per block. Each block started and ended with at least three standards. The first standard in each block was one of 12 possible orientation values, chosen randomly for each participant. For example, the first few trials of one sequence might be 8°… 8°… 8°… *68°* (i.e., +60° deviant for the first sequence …68°… 68°… 68°… 68°… *53°…* (i.e., −15° deviant for the second sequence). Figure 1(C) depicts these roving standard sequences.
2. Equiprobable blocks built from the roving standard blocks by replacing all its standard stimuli (outlined in green in Figure 1C)—except for the ones immediately preceding a deviant —with a Gabor patch whose orientation was randomly chosen among the 12 possible orientations Figure 1(D). All orientations appeared equally (8.3%) often. Deviants within oddball blocks (outlined in blue) are identical to the corresponding control stimulus (outlined in orange) in equiprobable blocks. The stimulus preceding a control stimulus (framed with two black chevrons) was identical to the stimulus preceding the deviant in the oddball block, so we can discount the possibility that a difference in pairing (in the oddball vs. control sequences) is contributing to any deviant-related differences.

There were six oddball blocks and six equiprobable blocks. We randomized block order afresh for each participant. Each block took less than four minutes to complete. Participants were free to take breaks between blocks.

### EEG recording and analysis

We recorded the electroencephalogram (EEG) using an EGI, 129-channel, dense-array HydroCel geodesic sensor net, and Net Amps 300 signal amplifier. We recorded EEG at a 500 Hz sampling rate. Impedances were below 50 kΩ as recommended by Ferree et al. [40] for the high-input impedance amplifiers. All electrodes were referenced to Cz.

We processed the EEG data offline using MATLAB 2015b (MathWorks Inc., USA), EEGLAB 14.1.1 [41], and ERPLAB 6.1.4 [42]. We re-referenced the signal of all electrodes to the common average and filtered the EEG with a low-pass 40 Hz Kaiser-windowed (beta 5.65) sinc finite impulse response (FIR) filter (order 184) followed by a high-pass 0.1 Hz Kaiser-windowed (beta 5.65) sinc FIR filter (order 9056). Epochs were 400 ms long, featuring a 50 ms pre-stimulus baseline to accommodate the short 360 ms stimulus-onset-asynchrony. We excluded epochs with amplitude changes exceeding 800 μV at any electrode

We identified electrodes with unusually high deviations in EEG activity relative to the average standard deviation pooled from all electrodes using the method described by Bigdely-Shamlo et al. [43]. A robust z-score was calculated for each electrode by replacing the mean by the median and the standard deviation by the robust standard deviation (0.7413 times the interquartile range). We removed any electrode with a z-score exceeding 2.0 provided at least four others surrounded them for later interpolation.

We performed independent component analysis (ICA) with AMICA [44]. To improve the decomposition, we performed the analysis on raw data (excluding bad electrodes) filtered by a 1 Hz high-pass (Kaiser-windowed sinc FIR filter, order 804, beta 5.65) and 40 Hz low-pass filter, segmented into epochs, but not baseline corrected [45] as suggested by Winkler et al. [46]. We simultaneously reduced the data to 32 components.

To ensure that trials where participants moved their eyes or blinked were not included in the final analysis of the data, we created vertical and horizontal EOG channels by bipolarizing data from electrodes above and below the right eye (electrodes 8 and 126) and outer canthi of both eyes (electrodes 1 and 32), respectively (as in [47]). We identified epochs containing amplitude changes exceeding ±60 μV at these EOG channels for rejection after ICA correction.

We applied the de-mixing matrix to the 0.1-40 Hz filtered data and used SASICA [48] to identify which components exhibited low autocorrelation, low focal electrode or trial activity, high correlation with vertical or horizontal EOG, or met ADJUST criteria [49]. We assessed the remaining components using criteria described by Chaumon et al. [50], classifying components based on consistent activity time-locked to stimulus onset across all trials, on topography, or on power spectrum. We removed components identified as unrelated to brain activity. We then removed epochs identified for rejection. We performed a final artifact rejection—removing epochs containing amplitude changes exceeding ±60 μV at any electrode. Finally, using spherical splines [51], we interpolated data for removed electrodes.

For exploring deviant-related responses, we averaged ERPs for the standard, deviant, and control trials, excluding epochs immediately following a deviant or control trial. We produced difference waves by subtracting ERPs to standards and ERPs to controls from ERPs to deviants. The mean number (SD) of epochs for standards were 1994 (261), 187 (26) for 15-degree deviants, 187 (28) for 15-degree controls, 186 (28) for 30-degree deviants, 188 (30) for 30-degree controls, 188 (25) for 60-degree deviants, and 188 (27) for 60-degree controls.

For exploring repetition suppression, we averaged ERPs for standard and random stimuli at varying positions, excluding deviant and control trials. We retained standard trials immediately following deviants—in oddball sequences—and random trials immediately following controls—in equiprobable sequences. The mean number (SD) of epochs for standards were 561 (76) for 2nd sequential position (SP), 568 (73) for 3rd SP, 473 (65) for 4th SP, 350 (47) for 5th SP, 239 (35) for 6th SP, 149 (23) for 7th SP, 90 (12) for 8th SP, 120 (21) for ≥9th SP. The mean number (SD) of epochs for randoms were 563 (89) for 2nd SP, 570 (86) for 3rd SP, 478 (73) for 4th SP, 351 (54) for 5 th SP, 239 (41) for 6th SP, 147 (26) for 7th SP, 90 (14) for 8th SP, and 123 (22) for ≥9th SP.

Using the EP Toolkit (v2.64; [19]), we conducted temporal principal component analysis (PCA) on the individual average ERP data for deviant and control trials from all 129 electrodes. We used Promax orthogonal rotation (*κ* = 3) with a covariance relationship matrix and Kaiser weighting as recommended by Dien et al. [52]. PCA reduced the data to those components explaining most of the observed data. The component that explained the greatest amount of signal is component 1 and every component thereafter explained less of the data than the component before it. We show how much of the signal a component is responsible for by plotting the component’s loading (scaled by SD) over time [53].

Using PCA, one can identify separate components in the ERP waveform and extract an alternative measure of ERP component amplitudes for inferential testing [52, 54, 55]. Each PCA component has a peak latency, a site of maximum positivity on the scalp (i.e., a component’s positive pole), and a site of maximum negativity on the scalp (i.e., a component’s negative pole). Plotting a topographical map of the microvolt scaled PCA data of a single component at the time of its peak latency shows the component’s positive and negative pole.

A vMMN component would emerge as a component that is largest (most negative) between 150-300 ms, based on previous vMMN studies using orientation deviants (e.g., 162-170 ms in [56]; 200-250 ms in [11]; 190-220 ms in [22]). A genuine vMMN component should also yield scores that are more negative for deviants compared to controls at the component’s negative pole.

We conducted Bayesian analyses of variance (ANOVAs) and paired *t*-tests to determine the likelihood of obtaining the data in addition to our traditional ANOVAs and paired *t*-tests. We used a medium prior (with a Cauchy prior whose width was set to 0.707) for all Bayesian analyses. For interpreting our findings, a model with the largest Bayes Factor (*BF*_10_) is the model that best explains the data; this is the favored model. All main effects and interactions in the favored model are, therefore, important for explaining the data. Evidence against the null is considered weak if a *BF*_10_ is between 1 and 3. It is positive for a *BF*_10_ between 3 and 20, strong for a *BF*_10_ between 20 and 150, and very strong given a *BF*_10_ greater than 150 [57].

We also performed Bayes Factor replication (*BF*_r0_) tests [58] on mean amplitudes between 197 and 207 ms at each PO ROI given a prior reflecting the effect size (Cohen’s *d* = −0.73) for the genuine vMMN at the PO8 in Kimura and Takeda’s [22] oddball condition for 32.7° orientation deviants.

For all frequentist analyses, we applied the Greenhouse-Geisser correction where necessary (ε<.750). Eta squared (η^2^) denotes the estimated effect size.

## Author contributions

Both authors contributed to the experiment’s conception, methodology, and writing – review & editing. AGM conducted formal analysis, data curation, investigation, project administration, visualization, and writing – original draft, review, and editing. ROS contributed to project administration and supervision.

## Acknowledgements and Funding

We are grateful to Urte Roeber, who contributed to conception, methodology, review, and supervision. We are also grateful to the students who assisted in collecting the data for this experiment. The research was supported by an Australian Government Research Training Program Scholarship to AGM.

## Supplementary Materials

### 1. Behavioral Results

Hit rates for oddball blocks (*M*=98.2%, *SD*=1.8%) were essentially identical for control blocks (*M*=98.2%, *SD*=2.3%), *t*(20) =0.016,*p*=.987, *BF*_10_=0.228. False alarm rates were similar in oddball (*M*=0.0007%, *SD*=0.0007%) and control blocks (*M*=0.0005%, *SD*=0.0005%), *t*(20)=1.678, *p*=.109, *BF*_10_=0.754. Reaction times were essentially identical in the oddball (*M*=545 ms, *SD*=52 ms) and control blocks (545 ms, *SD*=51 ms), *t*(20)=−0.025, *p*=.981, *BF*_10_=0.228.

### 2. Electrophysiological Results

#### 2.1. vNMN

Bayesian replication confirmed that there was neither a classic nor a genuine vMMN (complete analyses in Table S1). In fact, for the classic vMMN (green traces), there was a positivity in the analyzed time window—the opposite of a vMMN. Smout et al. showed the same positivity.

#### 2.2. Deviant-related positivity

For the deviant-related positivity (DRP), we performed a 3 × 3 × 2 repeated-measures ANOVA on P1 and N1 scores with parietal-occipital (PO) region (left vs. midline vs. right), magnitude of deviance (small vs. medium vs. large), and stimulus type (deviant vs. control) as factors.

For P1, magnitude of deviance was significant, *F*(2, 40)=26.811, *p*< .001, η^2^=.011, ε=.667. Holm post-hoc tests showed significantly larger scores for 60° orientation change compared with 15° (*t*=-7.282, *p*<.001, *d*=-0.280) and with 30° (*t*=-2.974, *p*=.005, *d*=-0.114) and for 30° compared with 15° (*t*=-4.308, *p*<.001, *d*=-0.166). Smout et al. found similar results, albeit in their attended condition.

A ROI × stimulus interaction emerged, *F*(2,40)=3.547, *p*=.038, η^**2**^<.001, ε=.907. Holm post-hoc tests showed deviants were significantly more positive than controls at LPO (*t*=2.882, *p*=.041, *d*=0.123), MPO (*t*=3.990, *p*=.003, *d*=4.293), and RPO (*t*=3.786, *p*=.010, *d*=3.786). The data provide very strong evidence for the favored Bayesian model including the main effects of magnitude of deviance, stimulus type, and ROI (*BF*_10_=3.275e +40).

There were no significant effects or interactions for the N1 although the data provide some positive evidence for the Bayesian model including the main effect of laterality only (*BF*_10_=15.961, *F*(2,40)=1.261, *p*=.294, η^2^=.037, ε**=**.928).

#### 2.3. Repetition suppression

To show the linear trends in scores for each stimulus, we show linear regressions in Figures 4 and S3 for P2 and N1 respectively. The P2 results show repetition suppression clearly, with the P2 scores declining with number of preceding standards and not with number of preceding random orientations. The N1 results are similar, except for the RPO data. Its two regression lines both show decreasing N1 scores of essentially identical slope. We must confess we have no idea why this weird result happened.

**Table S1.** Directed Paired (*BF*_10_) and Replication (*BF*_r0_) Bayesian t-tests of the Difference Wave Mean Amplitudes (μV) at Left (L), Middle (M), and Right (R) Parieto-occipital (PO) Regions Between 197 and 207 ms for Each Magnitude of Deviance (df = 20)

|  | Deviant vs. Standard (Classic) |  |  |  |  | Deviant vs. Control (Genuine) |  |  |  |  |
| --- | --- | --- | --- | --- | --- | --- | --- | --- | --- | --- |
| | $\mu V$ | $t$ | $p$ | $BF_{10}$ | $BF_{r0}$ | $\mu V$ | $t$ | $p$ | $BF_{10}$ | $BF_{r0}$ |
| *L PO* |  |  |  |  |  |  |  |  |  |  |
| 15° (small) | 0.41 | 3.879 | .999 | 0.061 | 0.009 | 0.18 | 1.516 | .927 | 0.101 | 0.012 |
| 30° (medium) | 0.55 | 7.701 | .999 | 0.016 | 0.014 | 0.04 | 0.294 | .614 | 0.185 | 0.038 |
| 60° (large) | 0.71 | 6.871 | .999 | 0.018 | 0.014 | 0.12 | 0.962 | .826 | 0.127 | 0.018 |
| *M PO* |  |  |  |  |  |  |  |  |  |  |
| 15° (small) | 0.36 | 3.471 | .998 | 0.064 | 0.008 | 0.16 | 1.060 | .849 | 0.122 | 0.017 |
| 30° (medium) | 0.69 | 4.597 | .999 | 0.056 | 0.009 | 0.25 | 1.465 | .921 | 0.103 | 0.012 |
| 60° (large) | 0.64 | 4.280 | .999 | 0.058 | 0.009 | 0.22 | 1.810 | .957 | 0.092 | 0.011 |
| *R PO* |  |  |  |  |  |  |  |  |  |  |
| 15° (small) | 0.17 | 1.933 | .966 | 0.088 | 0.010 | 0.06 | 0.387 | .649 | 0.174 | 0.033 |
| 30° (medium) | 0.48 | 3.464 | .999 | 0.064 | 0.008 | 0.25 | 1.892 | .963 | 0.089 | 0.010 |
| 60° (large) | 0.54 | 3.617 | .999 | 0.063 | 0.008 | 0.19 | 1.822 | .958 | 0.091 | 0.010 |

*Note.* Time-window is the same as the time-window in which Kimura and Takeda (2015) found the largest deviant-minus-control difference at PO8 for orientation deviants of 32.7°.

**Figure S1.**
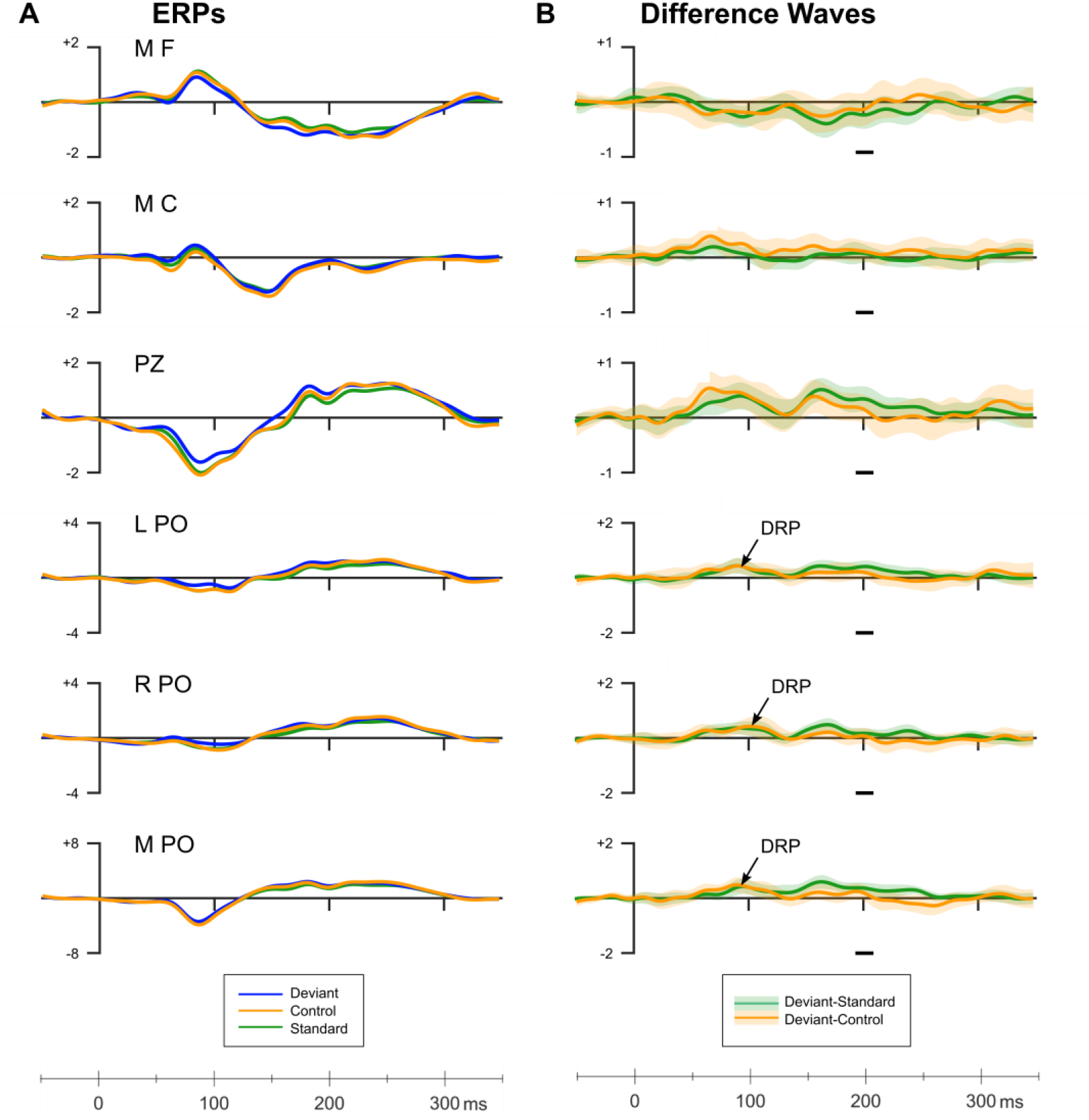
A. **Grand average ERPs for 15° orientation changes from various electrode clusters** around the electrodes Smout et al. reported. B. **Difference waves for classic (15-degree deviant minus standard) and genuine (15-degree deviant minus control) deviant-related activity**. The arrowed components show the only genuine deviant-related activity (DRP). Horizontal grey bars illustrate the time-window in which Kimura and Takeda (2015) found the largest deviant-minus-control difference (i.e., genuine vMMN) for 32.7° orientation deviants. Amplitudes from this time window were analyzed using Bayesian replication (results in Table S1). The lighter colors surrounding the difference waves give ± 1 standard error of the mean.

**Figure S2.**
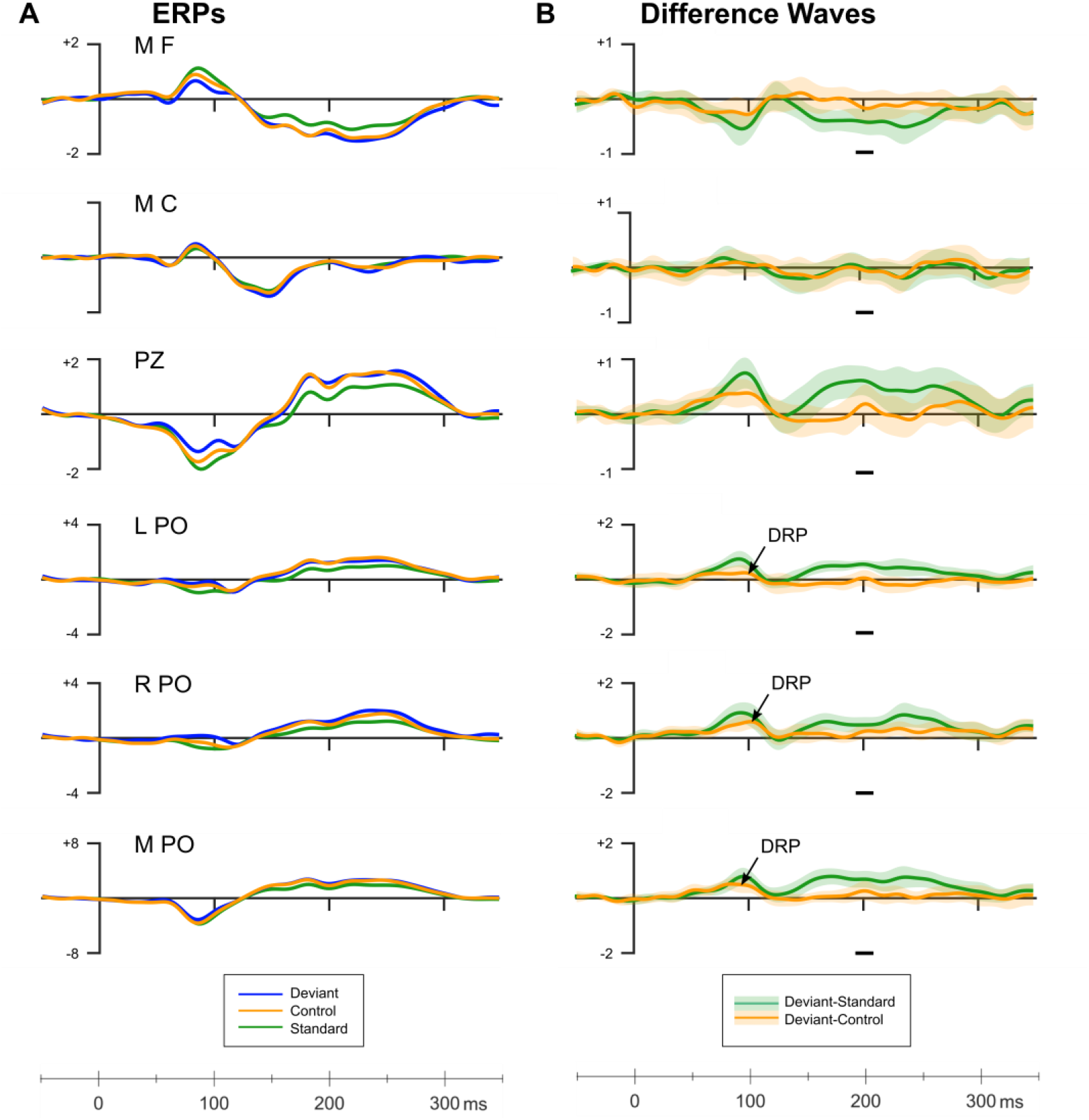
A. **Grand average ERPs for 30° orientation changes from various electrode clusters** around the electrodes Smout et al. reported. B. **Difference waves for classic (30-degree deviant minus standard) and genuine (30-degree deviant minus control) deviant-related activity**. The arrowed components show the only genuine deviant-related activity (DRP). Horizontal grey bars illustrate the time-window in which Kimura and Takeda (2015) found the largest deviant-minus-control difference (i.e., genuine vMMN) for 32.7° orientation deviants. Amplitudes from this time window were analyzed using Bayesian replication (results in Table S1). The lighter colors surrounding the difference waves give ± 1 standard error of the mean.

**Figure S3.**
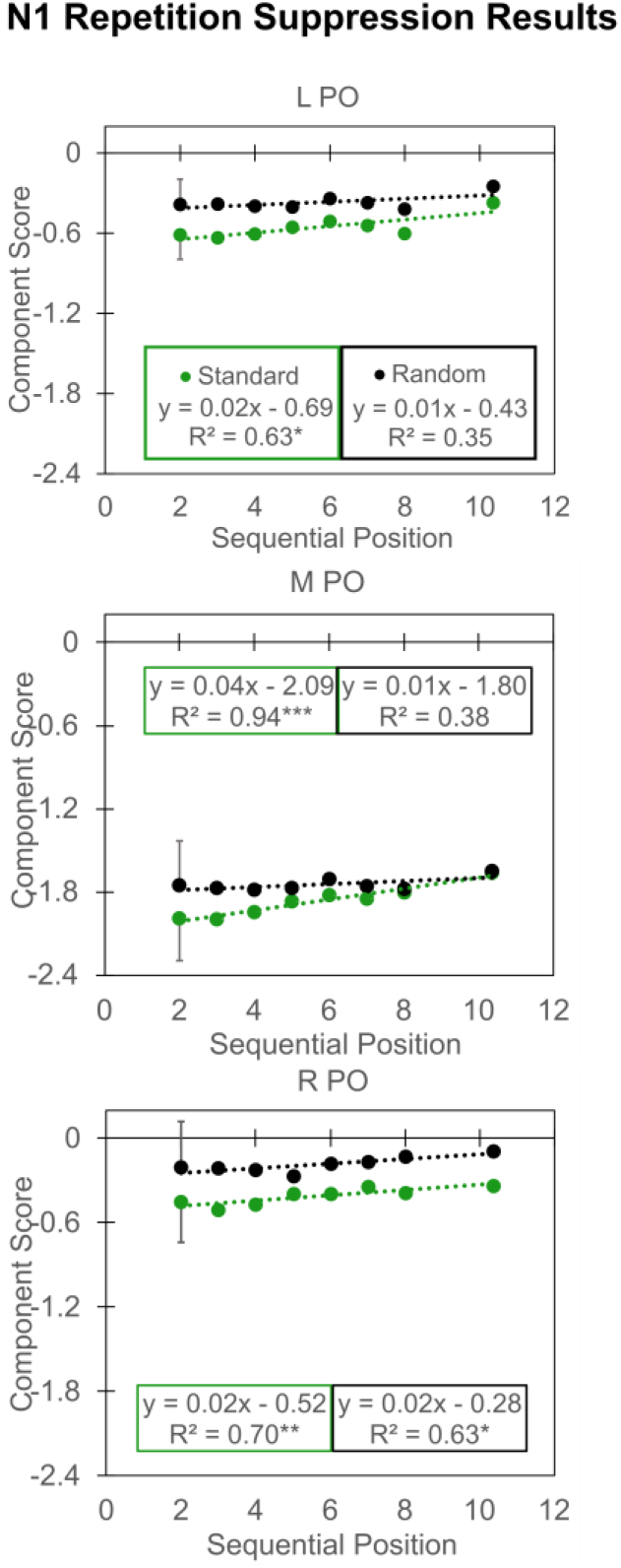
N1 repetition suppression results. N1 PCA scores are shown separately for the left (L), midline (M), and right (R) parietal-occipital (PO) regions of interest (ROI). Regression equations show the dotted lines in the data for standard and random stimuli. Asterisks denote the linear regression significance: \**p*<.05, \*\**p*<.01, *** *p*<.001. Error bars show 1 standard error (SE) calculated as the mean SE for all sequential positions. SE bars appear only on the second sequential position. We show +1 SE for standard and –1 SE for random stimuli.

